# Encoding Drug-Target-Pathway-Disease profiles for Drug-Cancer Association Prediction using Graph Transformer-Convolution Networks

**DOI:** 10.1101/2025.03.05.641572

**Authors:** Cheng Yu Tung, Jinn Moon Yang

## Abstract

New drug development is costly, time-consuming, and has a low success rate, leading to a decline in drug discovery efficiency over time. Drug repurposing has emerged as an effective alternative, applying existing, safe drugs to new diseases, thereby reducing development time and costs by bypassing preclinical toxicology testing. With the increasing availability of large-scale interaction data (e.g., drug-protein, protein-protein, and drug-disease networks) and advancements in generative AI, new opportunities have arisen for drug discovery. However, AI-based methods still face challenges: (1) ineffective integration of heterogeneous biological data across drugs, proteins, pathways, and diseases, and (2) lack of interpretability, limiting insights into drug mechanisms of action. To address these challenges, we propose a Graph Transformer-Convolution Network (GTCN) that integrates Graph Transformer Networks (GTNs) and Graph Convolution Networks (GCNs). By leveraging dynamic heterogeneous graph learning and attention mechanisms, our model optimizes relational structures within biological networks (drug-target-pathway-disease) and extracts more discriminative node features. Unlike traditional models that only encode direct drug-disease relationships, our approach captures how drugs act on proteins and regulate pathways to treat diseases. Furthermore, we design an interpretability framework that identifies critical elements for drug-cancer predictions, offering insights into disease mechanisms and drug mechanisms of action (MoA). This facilitates the discovery of new therapeutic strategies with biologically interpretable visualizations. The proposed dataset and code are available at https://github.com/zhengyutong99/GTCN

## Introduction

The development of new drugs is an arduous, time-consuming, and costly process, often exceeding 12–15 years with costs surpassing $2–3 billion USD [1]. Despite significant advances in computational modeling, high-throughput screening, and genomic sequencing, the efficiency of pharmaceutical research has declined, as described by “Eroom’s Law” [2]. The number of FDA-approved drugs per billion USD spent on research and development has halved approximately every nine years since 1950, highlighting the increasing complexity of drug discovery and regulatory requirements.

To address these inefficiencies, drug repurposing has emerged as an effective strategy, leveraging pre-existing data on approved drugs to identify new therapeutic applications [3]. By bypassing early-stage toxicology assessments, repurposing significantly reduces development time and cost while mitigating safety risks.

Currently, two major *in-silico* strategies are widely used in drug repurposing: target-based drug discovery (TDD) and phenotypic drug discovery (PDD) [4, 5]. Structure-based TDD focuses on identifying drug-target interactions based on molecular structures, making it effective for understanding disease mechanisms and designing precise inhibitors. However, its reliance on known targets limits its applicability to diseases with unclear molecular drivers [6, 7, 8]. In contrast, PDD identifies drugs based on cellular phenotypes rather than specific targets, making it useful for complex diseases. However, its “black box” nature complicates mechanistic interpretation and safety validation [4, 9, 10, 11].

With the increasing availability of large-scale biological data, such as drug-protein, protein-protein, and drug-disease interactions, deep learning has become a powerful tool for computational drug repurposing [12]. Graph-based approaches, particularly heterogeneous graph neural networks and attention-based models, have been widely explored to capture complex relationships within biological networks. Notable examples include MHGNN (Metapath-aggregated Heterogeneous Graph Neural Network), which encodes multi-relational biomedical data using meta-paths [13], and REDDA (Relations-Enhanced Drug-Disease Association), which introduces topological subnet embedding and graph attention mechanisms to improve drug-disease association prediction [14]. These models highlight the potential of heterogeneous graphs and attention mechanisms in drug repurposing. However, existing AI-driven approaches still face two major challenges:

- **Graph sparsity limits the utility of biological networks:** Current methods operate on highly sparse knowledge-base biological graphs, where many potential interactions are unrecorded or missing. As a result, they struggle to fully leverage the connectivity within biomedical data.
- **Lack of interpretability:** Many existing models operate as “black boxes,” offering limited insight into their decision-making processes. This makes it difficult for researchers and clinicians to trust the predictions, as the rationale behind the drug-disease associations remains unclear.

In this work, we propose a novel framework : **Graph Transformer-Convolutional Network (GTCN)**, which integrates **Graph Transformer Networks (GTNs)** [15, 16] **and Graph Convolutional Networks (GCNs)** [17]. While GTNs focus on editing and optimizing the network structure, GCNs process the refined network to extract meaningful features. Note that to address the issue of graph sparsity, we incorporate GTNs to refine the structure of biological networks. GTNs dynamically enhance or introduce biologically relevant interactions while weakening non-informative edge, effectively optimizing the connectivity within the network. This process improves the utilization of sparse biological graphs, allowing the model to extract more meaningful relationships from limited data.

In addition, while most existing methods only apply biologically meaningful encoding to the specific entities involved in prediction tasks (e.g., drug-disease association prediction encodes only drugs and diseases), our approach extends biologically meaningful encoding to all node types within the biological network. By incorporating biologically relevant features for **drugs, targets, pathways**, and **diseases**, our model fully leverages the network’s structural information, enabling a more comprehensive understanding of drug repurposing mechanisms.

Furthermore, we introduce an interpretability framework to enhance transparency in model predictions. In addition to analyzing attention scores to understand the importance of different edge types and their contribution, we also examine structural changes in the biological network during training. This allows us to interpret how the model identifies key interactions—such as which proteins are inhibited by a drug, which pathways are subsequently regulated, and how this leads to disease treatment. By incorporating this level of explainability, our model provides valuable insights into potential therapeutic mechanisms and offers a biologically interpretable framework for drug repurposing.

## Material and Methods

In this section, we provide a detailed description of the datasets utilized for constructing the GTCN network for Drug Repurposing, followed by a detailed explanation of the initial node representation and biological network encoding. We then provide the overall framework of GTCN.

### Dataset

Existing benchmarks in drug repurposing primarily focus on broad disease categories. In contrast, we propose a cell line-based benchmark, which incorporates four types of biological nodes—Drugs (D), Targets (T), Pathways (P), and Diseases (I)—along with six types of relationships (**Figure 1**). Specifically, we collect:

**Fig. 1.**
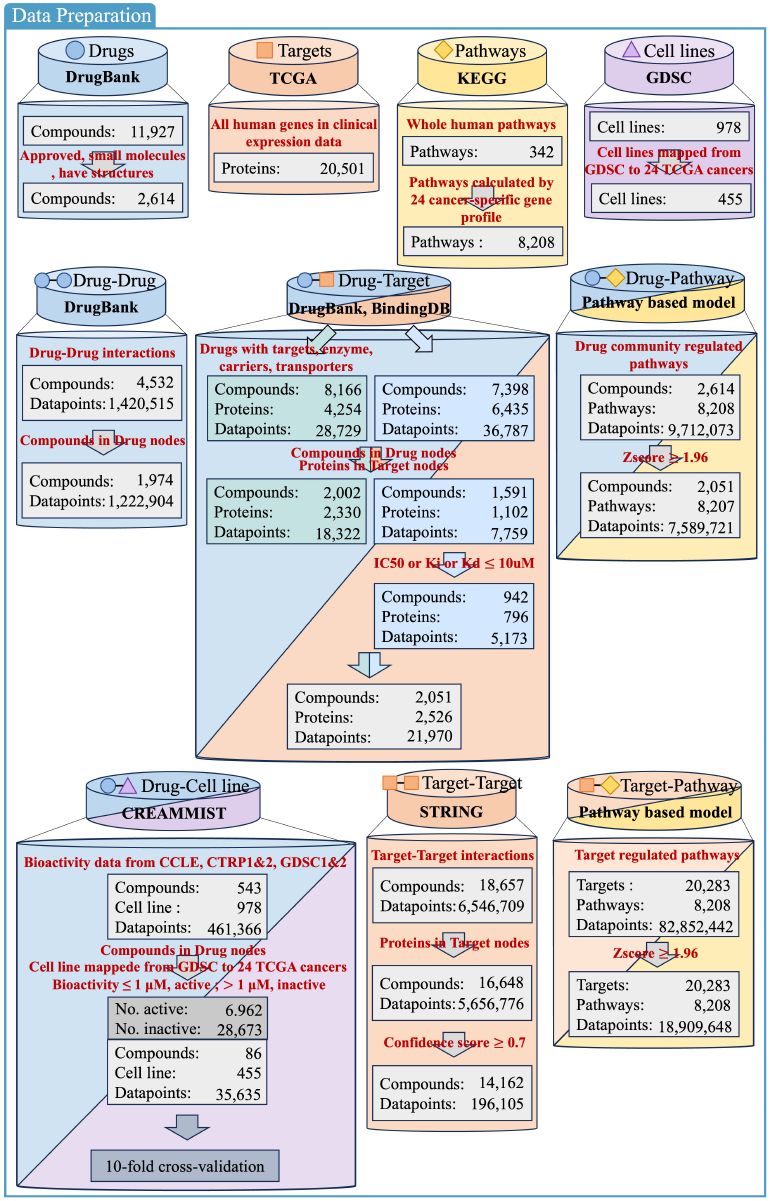
Data preparation

- **Drugs:** 2,614 approved small-molecule drugs from DrugBank (v5.1.11m) [18].
- **Targets:** 20,501 human proteins from TCGA [19].
- **Pathways:** 342 human pathways from KEGG [20]. Since we compute different drug-pathway (D-P) and target-pathway (T-P) relationships for 24 cancer types in TCGA, the total number of pathways expands to 8,208 (342 × 24).
- **Diseases:** 455 cancer-associated cell lines from GDSC [21], mapped to 24 cancer types.
- **Drug-Drug (D-D):** Drug interactions collected from DrugBank(v5.1.11m).
- **Drug-Target (D-T):**
  – Targets, enzymes, transporters, and carriers associated with drugs from DrugBank(v5.1.11m).
  – Protein targets from BindingDB, where drugs exhibit IC_50_, K_*i*_, or K_*d*_ ≤ 10μM [22, 23].
- **Drug-Pathway (D-P) and Target-Pathway (T-P):**
  – Computed using our in-house pathway-based model [24], which identifies pathways regulated by drug communities formed by single genes and drug targets.
- **Drug-Disease (D-I):** Drug treatment response in cell lines, where IC_50_ ≤ 10μM [25].
- **Target-Target (T-T):** Protein-protein interactions (PPIs) from STRING [26], filtered for confidence scores ≥ 0.7 [27, 28].

To emphasize the robustness and stability of GTCN, we also evaluate it on the benchmark dataset proposed by Gu et al. [14], which includes 894 drugs, 18,877 proteins, 20,561 genes, 314 pathways, and 454 diseases. This dataset consists of 10 types of biological relationships, covering drug-drug, drug-protein, protein-protein, protein-gene, gene-gene, gene-pathway, pathway-pathway, pathway-disease, disease-disease, and drug-disease interactions. The REDDA dataset is one of the few publicly available datasets that aligns closely with our research aim of integrating biological networks into a unified heterogeneous graph representation for learning.

### Node Representation

To effectively capture the biological characteristics of different entities in the network, we encode each node type with biologically meaningful features (**Figure 2**). This ensures that the model leverages domain-specific knowledge to enhance predictive performance.

**Fig. 2.**
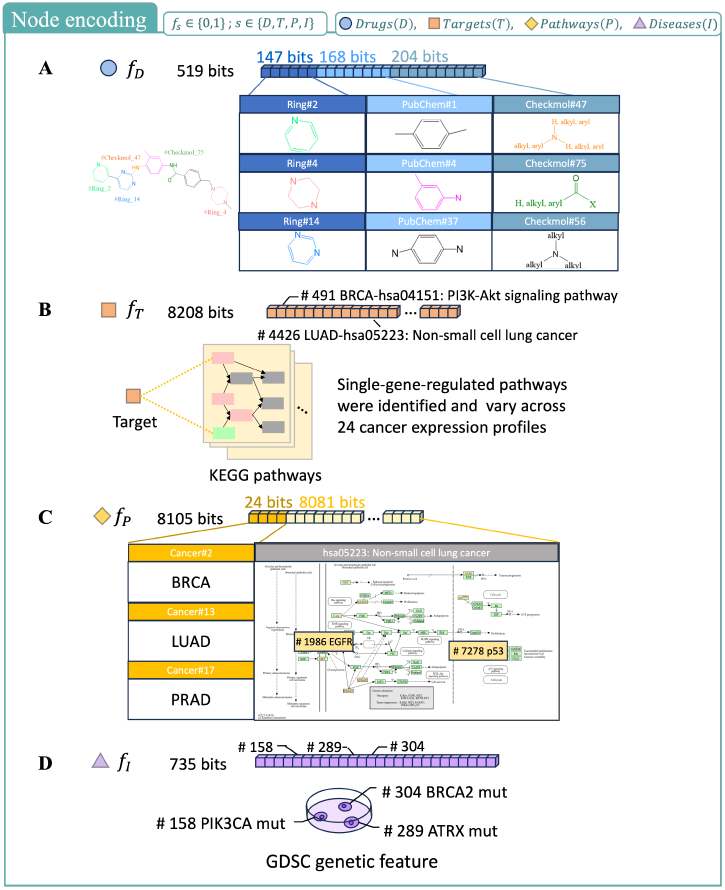
Node representation

For drug representation, we adopt a 519-bit structural fingerprint encoding, where each bit indicates the presence or absence of specific molecular moieties. These moieties are derived from CheckMol [29], PubChem [30], and Ring in Drug [31], allowing the encoding to capture key chemical properties relevant to drug interactions.

Protein targets are represented using an 8,208-bit pathway association vector, where each bit corresponds to a pathway that the protein significantly regulates. This encoding strategy incorporates functional pathway information, enabling the model to capture the biological role of each protein in disease mechanisms.

Pathway nodes are encoded based on their relevance to cancer-specific biological processes. Each pathway is assigned a feature representation consisting of 24 cancer-related attributes combined with 8,081 pathway-related protein interaction indicators, ensuring that the representation encapsulates both cancer relevance and regulatory protein involvement.

Finally, disease nodes, which correspond to cancer cell lines, are characterized by their genomic mutation profiles. Each disease is represented as a 735-bit binary mutation vector, where each bit denotes the presence or absence of a specific gene mutation. These mutations are extracted from GDSC, ensuring that the encoding accurately reflects disease-specific genetic variations.

### Adjacency Matrix Encoding

To effectively represent the heterogeneous biological network, we construct an adjacency matrix that encodes the relationships among drugs, targets, pathways, and diseases. Given the multi-relational nature of the dataset, we define a block-structured adjacency matrix where each block corresponds to a specific interaction type (**Figure 3**).

**Fig. 3.**
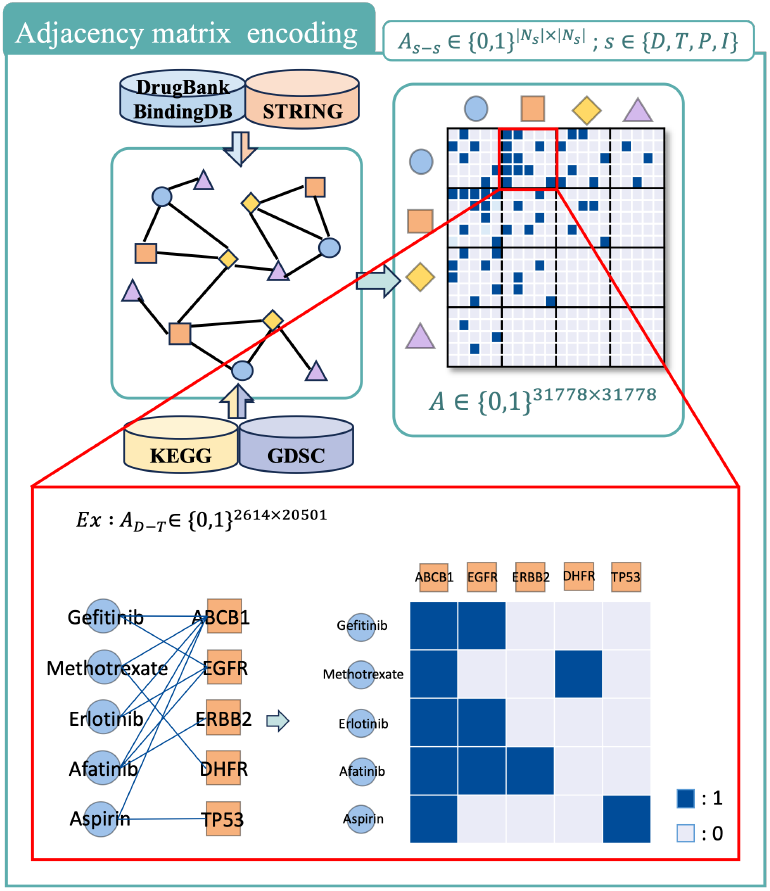
Adjacency matrix encoding

Each node type is assigned an index, and the adjacency matrix is partitioned accordingly. The D-D, D-T, D-P, T-P, D-I, and T-T interaction submatrices are extracted from their respective data sources. The adjacency matrix is composed of binary submatrices, each representing a specific edge type among two node categories. The size of a submatrix corresponds to the product of the number of nodes in the two categories involved. For instance, with 2,614 drugs and 20,501 targets, the Drug-Target (D-T) submatrix is of size 2, 614 × 20, 501. The six edge types collected form the primary submatrices. For asymmetric relationships (e.g., D-T or T-P), the transposed versions (T-D, P-D, I-D, P-T) are also included, resulting in a total of 10 submatrices. Each submatrix is indexed based on node indices for the corresponding types, ensuring consistency across the network.

### Feature Transformation

Subsequently, we applied a feature transformation matrix *W*_*s*_ specific to each node type, projecting the features into the same dimensional space:

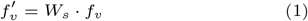

Here, 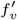 and *f*_*v*_ represent the transformed feature vector and the original feature vector of node *v*, respectively. The variable *s* ∈ {Drug, Target, Pathway, Disease} denotes the node type. The feature transformation addresses the heterogeneity of node features, facilitating better representation learning for downstream tasks.

### GTCN Architecture

The Graph Transformer-Convolutional Network (GTCN) is designed to address the limitations of existing drug repurposing models by integrating Graph Transformer Networks (GTNs) for dynamic network refinement and Graph Convolutional Networks (GCNs) for feature extraction (**Figure 4**). Unlike conventional approaches that rely on static biological networks, GTCN dynamically optimizes network topology which mitigates the sparsity issue commonly found in biological graphs. By leveraging GTNs to capture **global** structural dependencies and GCNs to perform **local** feature aggregation, GTCN ensures a comprehensive representation of multi-relational biological data. The learned node representations are subsequently used for predicting drug-cell line associations.

**Fig. 4.**
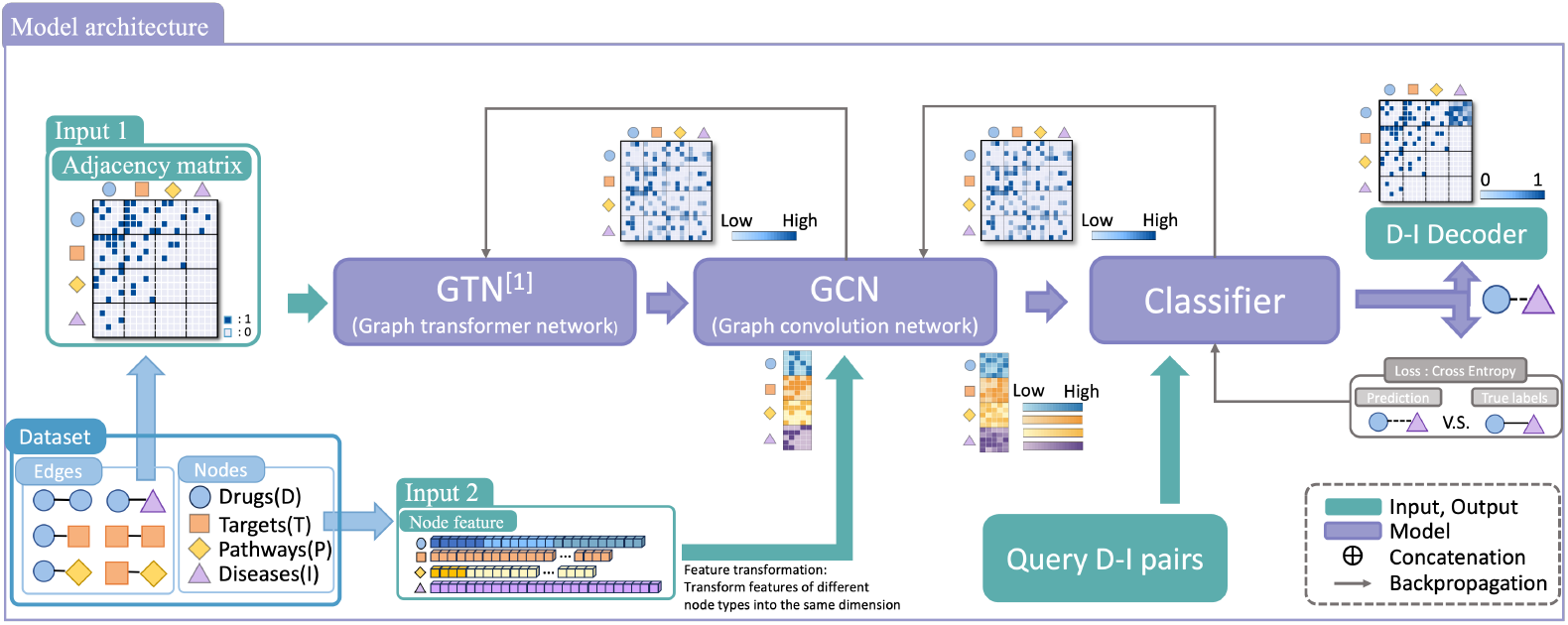
GTCN Architecture

#### Graph Transformer Networks

In the Graph Transformer layer, each submatrix is assigned a learnable attention score. These scores are used to weight the submatrices, emphasizing critical relationships while diminishing the impact of less important ones. The weighted submatrices are aggregated to produce the final heterogeneous adjacency matrix, where important connections are highlighted, and irrelevant ones are suppressed. By capturing **global** relational dependencies and dynamically modifying the network topology, GTNs enhance the overall network representation, ensuring that the model focuses on biologically meaningful relationships.

In GTNs, there are *m* Graph Transform Layers, where each layer learns weighted combinations of submatrices through a parameterized weight tensor. The weighted submatrices are generated via a 1 × 1 projection convolution operation, followed by a selection of suitable submatrix combinations. This process is formulated as:

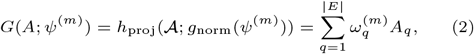

where *ψ*^(*m*)^ ∈ ℝ^1×1×|*E*|^ is a learnable parameter tensor that defines the importance of each edge type in the heterogeneous graph, with *E* representing the set of all edge types. The function *h*_proj_ applies a projection operation using 1 × 1 convolution, and 𝒜∈ ℝ^|*V* |×|*V* |×|*E*|^ represents the heterogeneous adjacency matrices of the node set *V*. Each adjacency matrix *A*_*q*_ corresponds to a specific edge type *q*, and the normalized weight 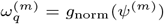 is obtained through a softmax normalization function.

The output adjacency matrices from each layer are aggregated into a comprehensive adjacency matrix, enabling all-to-all node connectivity. This integrated adjacency matrix is subsequently used in matrix multiplications to dynamically refine and adjust edge weights. To ensure numerical stability and prevent exploding values, the adjacency matrices undergo normalization using the following transformation:

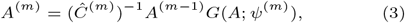

where *A*^(0)^ = *G*(*A*; *ψ*^(0)^) serves as the input adjacency matrix at the initial layer. The matrix *Ĉ*^(*m*)^ represents the degree matrix of *A*^(*m*)^, ensuring appropriate scaling during transformation.

#### Graph Convolutional Networks

After refining the heterogeneous graph with GTNs, we apply a Graph Convolutional Network (GCN) for feature learning. While GTNs focus on globally optimizing the network structure, GCNs emphasize **local** feature aggregation, ensuring that biologically meaningful node relationships are preserved. The GCN captures node interactions based on the refined adjacency matrix and propagates information across layers to learn expressive representations.

Given an initial node representation *Y* ^(0)^, set to 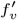 as defined in previous, node embeddings are updated layer by layer. For each GCN layer, the node representation *Y* ^(*l*)^ is updated according to:

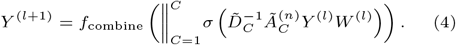

Here, *Y* ^(*l*)^ denotes the node embedding at layer *l*, while *C* represents the number of attention heads used in the transformation. The term 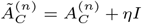 corresponds to the aggregated adjacency matrix of the *c*-th attention head after stacking *n* Graph Transform Layers, incorporating self-loops using *ηI* to ensure proper message passing. The degree matrix 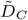 normalizes 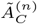 to mitigate potential numerical instability, and 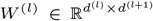 represents a trainable weight matrix. The function *f*_combine_ integrates the outputs from multiple attention heads.

#### Final Prediction and Optimization

The final node embeddings *Y* ^(*l*)^ are utilized for downstream drug-cell line association prediction. The embeddings are passed through fully connected layers (FC-Layers) followed by a softmax activation function:

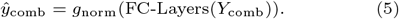

Here, *ŷ*_comb_ represents the predicted probability of drug-cell line associations, while *g*_norm_ applies a softmax function to normalize the output. The model is trained using a cross-entropy loss function:

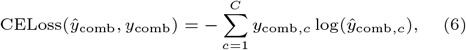

where *y*_comb,*c*_ is the ground truth label (either 0 or 1), and *C* denotes the number of classes. The parameters of the model are optimized using the Adam optimizer to minimize the cross-entropy loss, ensuring robust convergence during training.

#### Experimental and Hyper-parameter Settings

We conduct a 10-fold cross-validation on both our dataset and the benchmark dataset proposed by Gu et al. [14]. To evaluate the performance of GTCN in comparison with other models, we adopt multiple evaluation metrics, including AUC, F1-score, Accuracy, Recall, Specificity, Precision, and MCC, which are computed following the process outlined in Ref. [32]. Notably, the performance of other models in our comparisons is directly obtained from the results reported in Gu et al.’s published work.

Regarding the hyper-parameter settings, due to GPU memory constraints, we utilize a single Graph Transform Network (GTN) layer and three Graph Convolutional Network (GCN) layers. The Adam optimizer is employed with a learning rate ranging from 10^−3^ to 10^−5^, and the maximum number of epochs is set to 5000. However, early stopping is applied if the loss does not continuously converge during training.

## Result

### Performance Comparison with state-of-the-art approaches on the REDDA Dataset

### Performance Evaluation on REDDA Benchmark

#### Dataset

In this study, we compared the performance of our proposed GTCN against other methods on the REDDA dataset, these methods including SCMFDD [33], MBiRW [34], DDA-SKF [35], NFigureCN [**?**], HINGRL-Node2Vec-RF, HINGRL-DeepWalk-RF [36], LAGCN [37], DRWBNCF [38], and REDDA (**Figure 5A**). GTCN demonstrated superior performance in identifying positive samples (drug-disease associations), as reflected by its higher recall, precision, and F1-score. This indicates the effectiveness of our model in capturing meaningful biological associations that are often overlooked by other approaches. However, despite our model’s strengths, it showed marginally lower performance in accuracy and specificity compared to certain methods. A deeper examination of the REDDA dataset revealed that **99.93% of the labels are negative samples**, leaving only **0.07% as positive samples (Figure 5B)**. This extreme class imbalance heavily biases models towards predicting negative relationships, allowing methods that rely on simplistic assumptions (e.g., always predicting no drug-disease relationship) to achieve artificially high accuracy and specificity. For instance, methods leveraging this imbalance achieve high performance by merely classifying all samples as negatives, thereby neglecting the critical task of identifying true drug-disease associations.

**Fig. 5.**
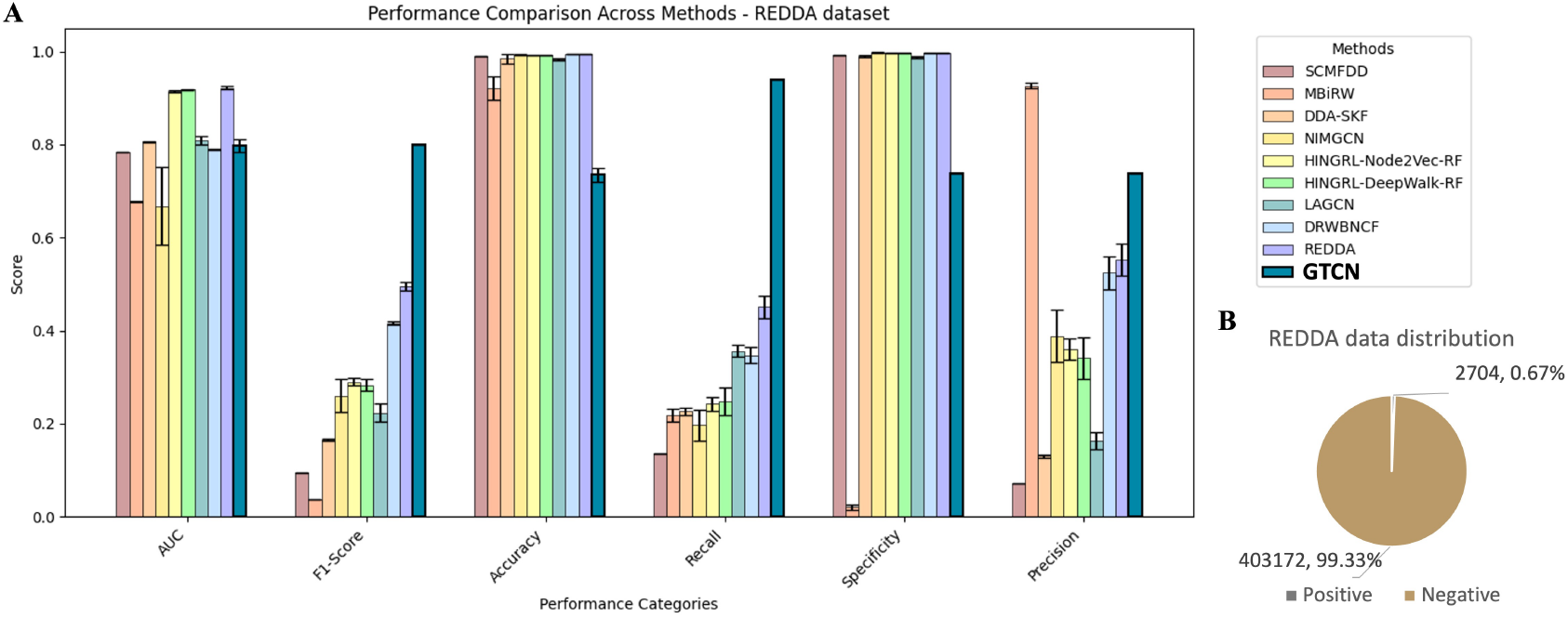
Performance Evaluation on REDDA Benchmark Dataset. (A) Performance comparison of our model with other drug-diseases association prediction methods on the REDDA dataset across various metrics. (B) Data distribution in the REDDA dataset showing the severe imbalance, with 99.33% of the labels as negative samples and only 0.67% as positive samples.

### Performance Evaluation on Our Proposed Cell Line-Based Dataset

To further enhance drug repurposing research, we propose a cell line-based drug-disease dataset as an alternative benchmark for model evaluation. By incorporating biological insights from cancer cell lines, our dataset provides a more refined framework for assessing drug-disease associations. As shown in **Figure 6A**, the dataset maintains a reasonable distribution between positive and negative samples, ensuring reliable model assessment.

**Fig. 6.**
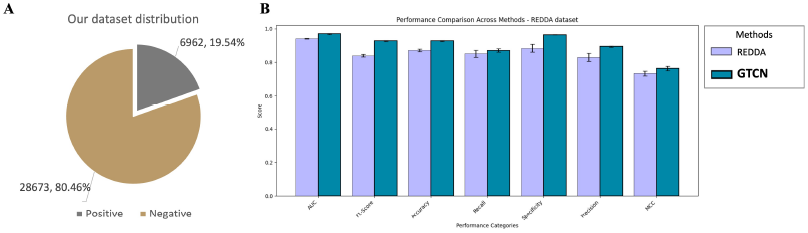
Performance Evaluation on Our Cell Line-Based Dataset. (A) Data distribution in our cell line-based dataset. (B) Comparison of performance metrics between the REDDA model and GTCN on our dataset. GTCN demonstrates superior performance across all metrics, indicating its robustness and effectiveness in drug-disease association prediction.

We benchmarked both the REDDA model and our GTCN on this dataset, and the results (**Figure 6B**) demonstrate that our model consistently outperforms REDDA across all key metrics. This superior performance highlights GTCN’s ability to capture biologically meaningful associations and generalize across complex biomedical networks. These findings not only validate the effectiveness of our model but also emphasize the importance of using biologically contextualized datasets to improve the reliability of computational drug repurposing approaches.

### Effect of Encoding Methods

#### Expanding Biological Context Through Node Encoding

Current approaches for predicting biological relationships often encode only the node types directly involved in the prediction task, overlooking the broader biological context. For instance, [14] only assigned meaningful encodings to drug and disease nodes, while proteins, genes, and pathways were left as zero vectors. Similarly, [13] employed one-hot encodings for all nodes. In contrast, we propose encoding all biological node types to enhance the model’s ability to capture intricate interactions within complex networks. A comparison of different encoding methods is provided in **Table 1**.

**Table 1.** Comparison of Node Encoding Methods

| Node | REDDA Method | Our Method |
| --- | --- | --- |
| Drug | Drug-Drug similarity | Moiety-based encoding |
| Target | Zero vector | Single-gene regulated pathways |
| Pathway | Zero vector | Protein-based pathway encoding |
| Gene | Zero vector | Not encoded |
| Disease | Disease-Disease similarity | Gene mutation profiles |

#### Experimental Evaluation of Encoding Methods

To assess the impact of encoding strategies, we applied our biologically meaningful encoding to the REDDA model and compared it to the original REDDA encoding. As shown in **Figure 7**, our method significantly outperforms the baseline across all metrics, particularly in imbalanced datasets, where it improves MCC by 55%. This highlights the importance of encoding biological context in learning meaningful relationships.

**Fig. 7.**
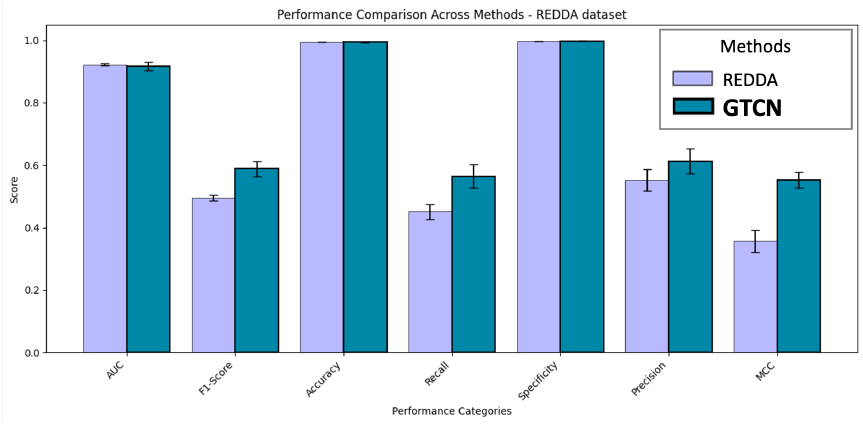
Performance Comparison Using Different Encoding Methods. Comparison of the REDDA model’s performance using its original encoding and our biologically meaningful encoding. Metrics include AUC, F1-score, accuracy, recall, specificity, precision, and MCC.

We further compared our biologically enriched encodings with one-hot encoding ([13]) across various node types using GTCN and our dataset. We started with all nodes initialized as one-hot vectors and incrementally replaced them—first encoding drugs using moiety-based features, then incorporating disease-related gene mutation profiles, followed by pathway and target encodings. As summarized in **Table 2**, replacing one-hot encodings with biologically meaningful representations consistently improved model performance across all metrics. The most significant gains were observed when drug and disease encodings were introduced, particularly in recall and MCC, highlighting their crucial role in capturing meaningful biological interactions. Further improvements were achieved as pathway and target nodes were also assigned biologically relevant features, ultimately yielding the best overall performance. This stepwise enhancement suggests that each biologically encoded node type uniquely contributes to model effectiveness, reinforcing the importance of leveraging domain knowledge in graph-based drug repurposing models.

**Table 2.** Comparison of Encoding Strategies. Performance comparison of our model under different encoding strategies. Results are shown for one-hot encoding and biologically meaningful encodings applied to drug, target, pathway, and disease nodes. The best-performing value for each metric is highlighted in **bold**, and the lowest-performing value is <u>underlined</u>.

| Encoding Methods | AUC | F1-Score | Accuracy | Recall | Specificity | Precision | MCC |
| --- | --- | --- | --- | --- | --- | --- | --- |
| One-hot | 0.9451 ± 0.0150 | 0.8439 ± 0.0234 | 0.9039 ± 0.0234 | 0.8202 ± 0.0250 | 0.9600 ± 0.0649 | 0.8633 ± 0.0171 | 0.7331 ± 0.0242 |
| Drug | 0.9686 ± 0.0029 | 0.8807 ± 0.0075 | 0.9267 ± 0.0046 | 0.8690 ± 0.0078 | 0.9635 ± 0.0035 | 0.8924 ± 0.0081 | 0.7609 ± 0.0150 |
| Drug, Disease | **0.9692 ± 0.0029** | 0.8803 ± 0.0085 | 0.9269 ± 0.0042 | 0.8682 ± 0.0098 | 0.9646 ± 0.0020 | 0.8934 ± 0.0056 | 0.7611 ± 0.0148 |
| Drug, Disease, Pathway | 0.9689 ± 0.0029 | 0.8796 ± 0.0074 | 0.9265 ± 0.0043 | 0.8687 ± 0.0091 | 0.9645 ± 0.0037 | 0.8920 ± 0.0070 | 0.7603 ± 0.0145 |
| Drug, Disease, Pathway, Target | 0.9690 ± 0.0028 | **0.8808 ± 0.0074** | **0.9275 ± 0.0043** | **0.8687 ± 0.0084** | **0.9652 ± 0.0030** | **0.8946 ± 0.0072** | **0.7629 ± 0.0146** |

Our results confirm that biologically meaningful encodings improve model performance across all evaluation metrics. By capturing rich biological relationships, our approach enhances predictive accuracy and provides a more robust framework for computational drug repurposing.

### Effect of Graph Transformer Networks

Biological networks are highly incomplete, with knowledge-based networks covering only 10% of potential edges. To enhance connectivity, we incorporated Graph Transformer Networks (GTNs) into our model for drug-cancer association prediction. GTNs refine existing connections by strengthening critical links, weakening less relevant ones, and introducing new informative edges, enriching the biological network.

To assess GTNs’ impact, we compared model performance with and without the module. As shown in **Figure 8**, integrating GTNs significantly improves specificity and MCC, demonstrating enhanced ability to capture meaningful biological relationships. The MCC increase particularly highlights GTNs’ effectiveness in handling imbalanced datasets. These results confirm the importance of GTNs in refining biological networks, leading to more accurate predictions.

**Fig. 8.**
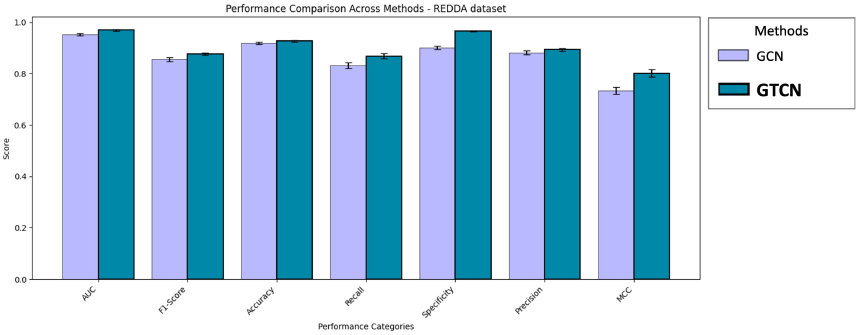
Performance Comparison With and Without GTNs. This figure compares the performance of the model using only GCN versus the full architecture with GTNs. Performance improves when GTNs are integrated, highlighting the module’s role in refining biological networks.

### Ablation Study: Identifying Optimal Parameters

To refine our model’s design, we conducted an ablation study to assess the impact of key hyperparameters, including the number of attention heads in GTNs and the classifier input dimension.

#### Impact of Attention Head Number

GTNs assign attention scores to submatrices of the adjacency matrix, influencing how connections are refined. We evaluated configurations with 1, 2, 3, and 4 attention heads (**Table 3**). While performance variations were minor, three heads achieved the best overall results, balancing computational efficiency and network refinement. These findings suggest that while head count does not drastically affect performance, optimizing it can enhance resource utilization.

**Table 3.** Impact of Attention Head Number on Performance. Performance of the model with different numbers of attention heads. The best-performing value for each metric is highlighted in **bold**, while the lowest is <u>underlined</u>.

| Heads | AUC | F1-Score | Accuracy | Recall | Specificity | Precision | MCC |
| --- | --- | --- | --- | --- | --- | --- | --- |
| 1 | 0.9686 ± 0.0027 | 0.8807±0.0063 | 0.9275 ± 0.0036 | 0.8684 ± 0.0077 | 0.9654 ± 0.0022 | 0.8949 ± 0.0054 | 0.7628 ± 0.0123 |
| 2 | 0.9688 ± 0.0031 | 0.8796±0.0090 | 0.9270 ± 0.0051 | 0.8667 ± 0.0110 | 0.9656 ± 0.0036 | 0.8946 ± 0.0083 | 0.7607 ± 0.0177 |
| 3 | **0.9690 ± 0.0028** | **0.8828±0.0074** | **0.9276 ± 0.0043** | **0.8687 ± 0.0084** | <u>0.9652 ± 0.0030</u> | 0.8946 ± 0.0072 | **0.7629 ± 0.0146** |
| 4 | 0.9686 ± 0.0028 | 0.8807±0.0080 | 0.9276 ± 0.0047 | 0.8679 ± 0.0089 | **0.9659 ± 0.0032** | **0.8956 ± 0.0079** | 0.7629 ± 0.0159 |

#### Impact of Classifier Input Dimension

We analyzed the effect of classifier input dimensions on performance (**Table 4**). Specifically, the output embeddings of drugs and diseases from the GCN have dimensions of 32, 64, 128, 256, or 512, respectively. Before being passed into the classifier, these embeddings are concatenated, resulting in final input dimensions of 64 (32+32), 128 (64+64), 256 (128+128), 512 (256+256), and 1024 (512+512). Our results indicate that an input dimension of 128 (64+64) achieves the best balance between information richness and computational efficiency. Larger dimensions introduce redundancy, leading to performance decline, while smaller dimensions may not capture sufficient biological context.

**Table 4.** Impact of Classifier Input Dimension on Performance. Performance of the model with different input dimensions. The best-performing value for each metric is highlighted in **bold**, while the lowest is <u>underlined</u>.

| Dimension | AUC | F1-Score | Accuracy | Recall | Specificity | Precision | MCC |
| --- | --- | --- | --- | --- | --- | --- | --- |
| 32 | 0.9614 ± 0.0032 | 0.8801 ± 0.0078 | **0.9270 ± 0.0044** | 0.8633 ± 0.0098 | **0.9647 ± 0.0021** | **0.8936 ± 0.0061** | 0.7604 ± 0.0153 |
| 64 | **0.9692 ± 0.0029** | **0.8803 ± 0.0085** | 0.9269 ± 0.0042 | **0.8682 ± 0.0098** | 0.9646 ± 0.0020 | 0.8934 ± 0.0056 | **0.7611 ± 0.0148** |
| 128 | 0.9588 ± 0.0028 | 0.8700 ± 0.0076 | 0.9168 ± 0.0044 | 0.8592 ± 0.0090 | 0.9537 ± 0.0028 | 0.8822 ± 0.0068 | 0.7611 ± 0.0149 |
| 256 | 0.9587 ± 0.0030 | 0.8696 ± 0.0073 | 0.9166 ± 0.0043 | 0.8580 ± 0.0083 | 0.9543 ± 0.0031 | 0.8828 ± 0.0072 | 0.7503 ± 0.0144 |
| 512 | 0.9487 ± 0.0028 | 0.8588 ± 0.0070 | 0.9164 ± 0.0040 | 0.8463 ± 0.0086 | 0.9541 ± 0.0020 | 0.8732 ± 0.0057 | 0.7490 ± 0.0137 |

This study optimized hyperparameter selection, enhancing model efficiency while maintaining predictive power. Due to hardware limitations, GTNs were restricted to two layers and GCNs to three. Future work will explore deeper architectures and more efficient adjacency matrix encoding strategies to further improve performance while addressing computational constraints.

### Model Interpretability: Observation of Attention Scores

#### Effect of Attention Score Initialization Methods

To enhance model interpretability, we analyzed the evolution of attention scores in Graph Transformer Networks (GTNs). These scores highlight the importance of different edge types in prediction. Initially, we used random initialization with softmax normalization; however, we observed that certain edge types failed to converge properly, with their attention scores still changing significantly at the final epoch. To address this issue, we introduced equal initialization across all edge types, enforcing a uniform starting point for attention scores (**Figure 9**).

**Fig. 9.**
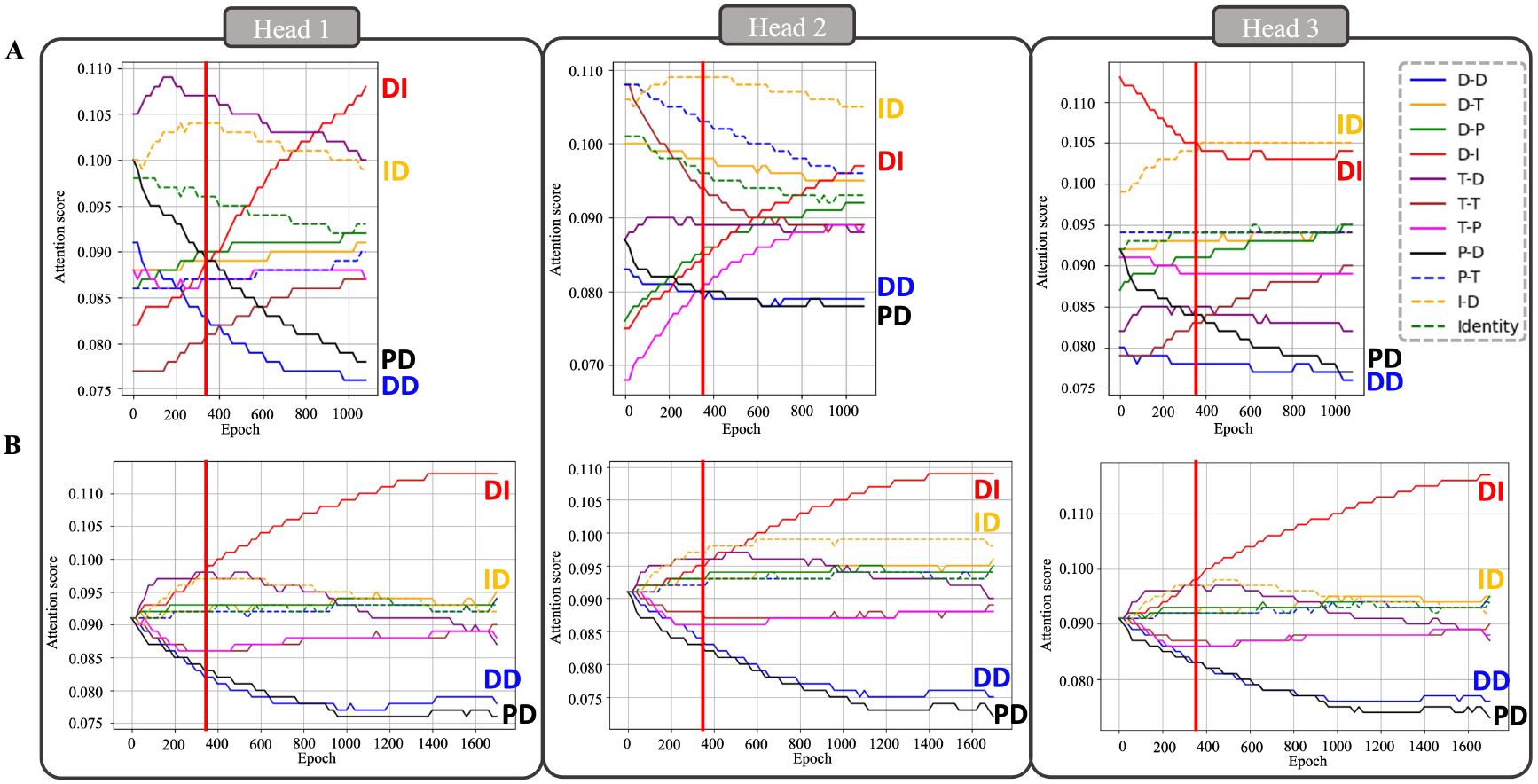
Attention Score Trends Across Edge Types. (A) Randomly initialized scores with standard softmax normalization. (B) Equally initialized scores across all edge types. Red vertical lines mark the loss convergence epoch.

This modification led to more stable and interpretable attention trends, with Drug-Disease (D-I) and Disease-Drug (I-D) edges consistently receiving the highest weights—aligning with the model’s primary task. Conversely, Drug-Drug (D-D) and Pathway-Disease (P-D) edges became less significant over time. Without equal initialization, some edges (e.g., D-I in Head 2) struggled to reach their expected importance levels by the end of training, highlighting the benefits of balanced initialization in ensuring robust convergence.

Further analysis (**Figure 10**) reveals that model convergence is determined based on the stabilization of training and testing loss, occurring around 350 epochs. Before this point, attention scores exhibit significant fluctuations as the model explores the relative importance of different edge types. After convergence, key edges such as D-I stabilize at higher values, while less critical edges (e.g., P-D) continue to decline. These findings align with existing studies emphasizing the importance of early-stage exploration and late-stage refinement in attention mechanisms. The persistent adjustments to critical edges, such as D-I, highlight their pivotal role in learning drug-disease associations. Conversely, the diminishing importance of less relevant edges suggests that the model effectively filters out redundant or non-contributory information.

**Fig. 10.**
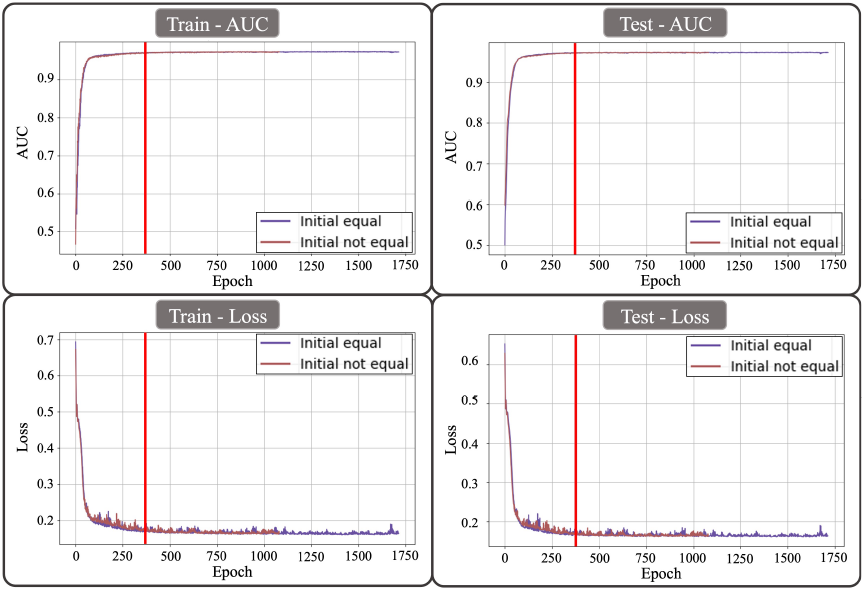
Model Convergence During Training. Red vertical line at epoch 350 marks convergence. Top row: AUC trends for training/testing sets. Bottom row: Loss trends for training/testing sets.

#### Effect of Drug-Drug Encoding on Attention Scores

We further examined how different D-D encoding strategies impact attention scores (**Figure 11**): (A) DrugBank-recorded interactions, (B) continuous Tanimoto similarity scores, and (C) top-15 binarized Tanimoto similarity. Across all methods, D-I remained the most weighted edge, while P-D had the lowest weight. Notably, similarity-based encodings (B, C) increased the importance of D-D and T-D edges, suggesting they introduce additional discriminative information.

**Fig. 11.**
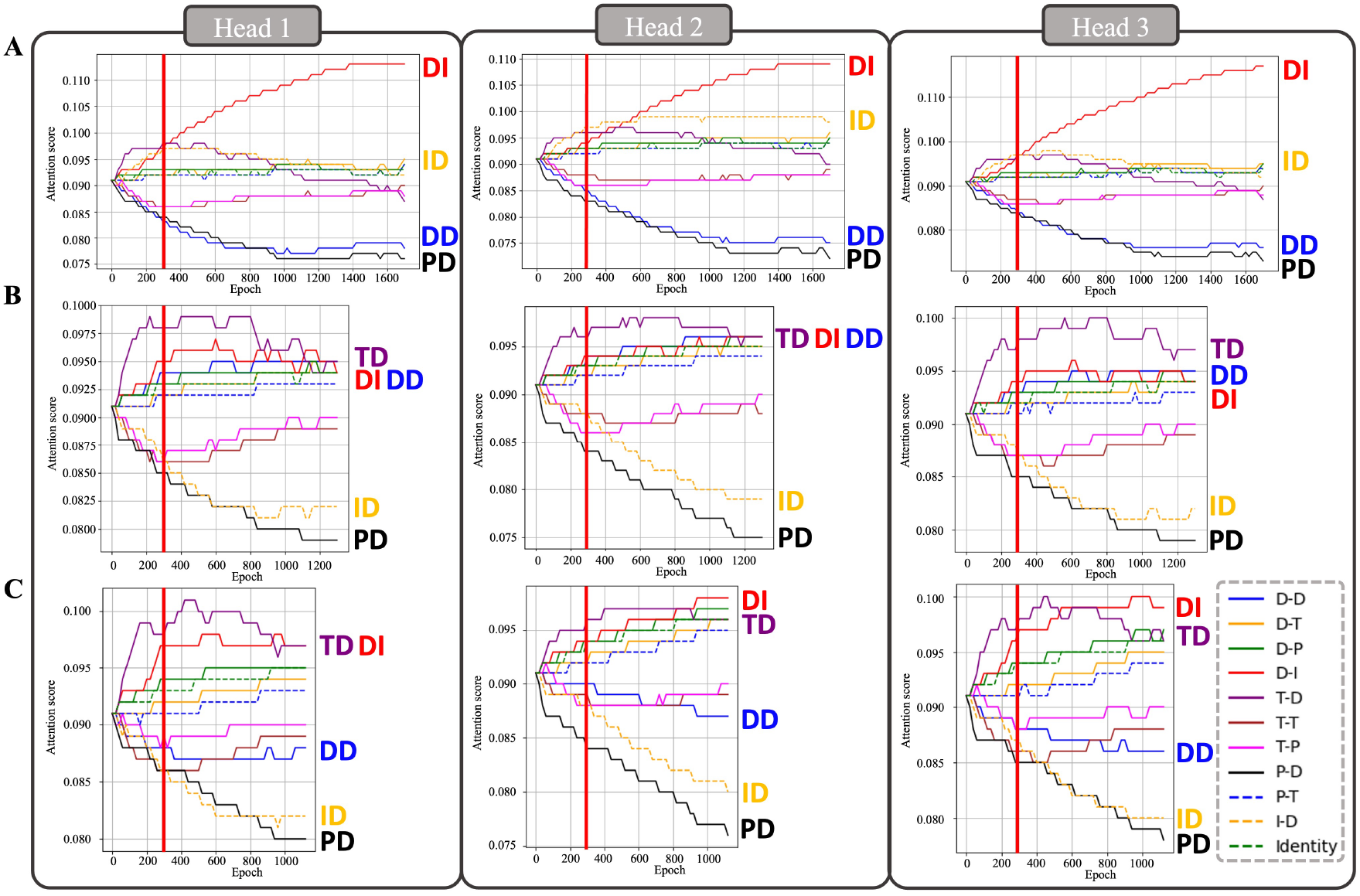
Attention Score Trends with Different D-D Encodings. (A) DrugBank-based edges. (B) Tanimoto similarity (continuous). (C) Tanimoto similarity (binary, top 15). Red vertical lines mark the convergence point (epoch 300).

The convergence analysis (**Figure 12**) shows that all models stabilize loss and attention scores around 300 epochs, though critical edges like D-I and I-D continue refining beyond this point. This suggests that similarity-based encodings help the model adaptively prioritize biologically meaningful relationships.

**Fig. 12.**
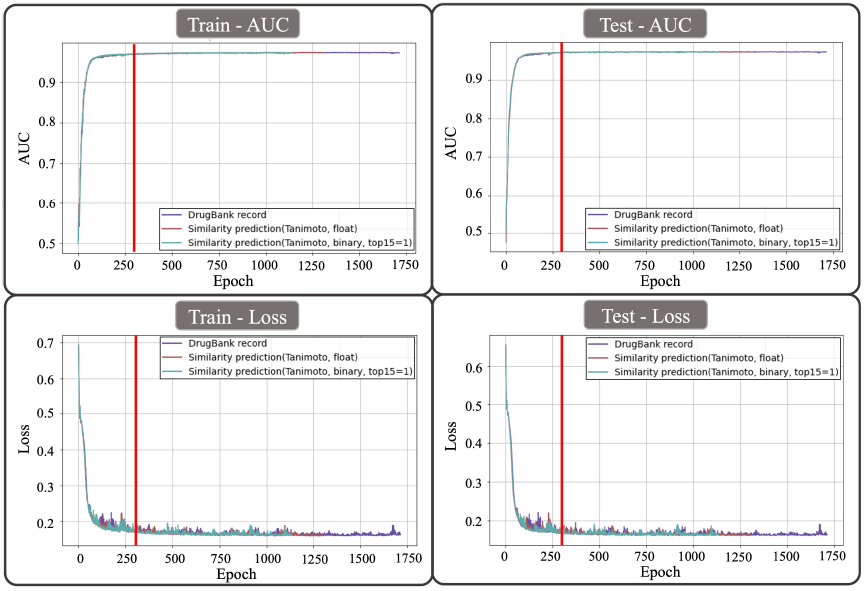
Model Convergence with Different D-D Encodings. Red vertical line at epoch 300 marks convergence. Top row: AUC trends for training/testing sets. Bottom row: Loss trends for training/testing sets.

### Case Study: Interpreting Model Decisions through D-T-P-I Relationships

To understand how our model refines biological networks for drug-disease prediction, we examined changes in drug-target-pathway-disease (D-T-P-I) relationships across training epochs. Since drug efficacy is driven by the inhibition of specific protein targets, leading to pathway regulation and subsequent disease treatment, analyzing the evolution of D-T-P-I connections provides insights into how the model captures these mechanisms. By observing how the model dynamically adjusts these relationships, we can better interpret its decision-making process.

To demonstrate the generalizability of our Graph Transformer-Convolutional Network (GTCN), we selected two case studies: Methotrexate, a chemotherapy drug, and Temsirolimus, a targeted therapy. These cases highlight that our model is not restricted to specific drug types but can effectively capture key biological interactions across different therapeutic modalities. Given the complexity of biological networks, we focused on the top-ranking relationships identified by the model, as these are most likely to contribute to the predictions.

**Figures 13** and **14** illustrate these case studies, where different colored boxes represent key changes in the network across training epochs. Blue boxes indicate nodes that were initially ranked high but were removed by the final epoch, while green boxes denote newly recognized key nodes. Red boxes highlight literature-supported pathways and diseases, while purple boxes represent potential drug repurposing opportunities. Orange and blue boxes with numbers indicate the ranking of drug-target relationships, while orange-yellow boxes indicate target-pathway rankings. This visualization helps reveal how the model prioritizes biological interactions during training.

**Fig. 13.**
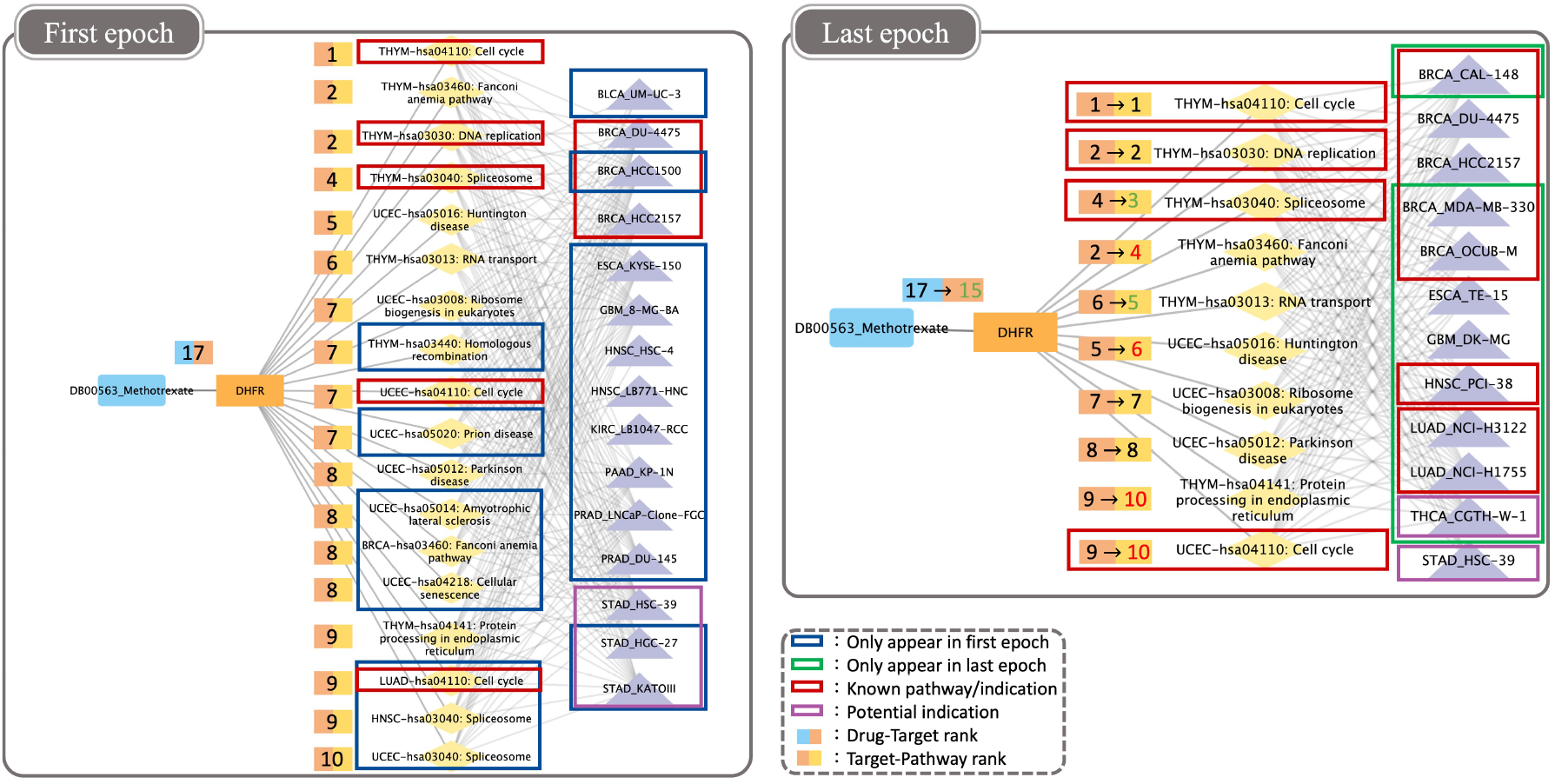
Case Study 1: D-T-P-I Networks of Methotrexate.

#### Methotrexate Case Study

For the chemotherapy drug Methotrexate (**Figure 13**), the model identified its known target DHFR [39] as a top-ranking target early in training. The known treatment relationship with breast cancer (BRCA) [40] and the potential treatment relationship with stomach adenocarcinoma (STAD) [41] were preserved throughout training. Additionally, the model retained key cancer-related pathways, such as those involved in cell cycle regulation, DNA replication, and RNA processing (spliceosome, RNA transport) [42], all of which are well-documented mechanisms affected by Methotrexate.

Beyond preserving these known interactions, GTCN progressively refined its predictions and identified additional established indications, including head and neck squamous cell carcinoma (HNSC) and lung adenocarcinoma (LUAD), both of which are already recognized as Methotrexate treatment targets [43, 44]. Notably, the model also suggested a new potential treatment link between Methotrexate and thyroid cancer (THCA), aligning with preclinical evidence in animal models [45].

#### Temsirolimus Case Study

For the targeted therapy drug Temsirolimus (**Figure 14**), the model successfully identified its known target mTOR [46], as a top-ranking target. Initially, renal cancer (KIRC), a known indication of Temsirolimus, was not highly ranked but became prominent by the final epoch [47]. The model also highlighted pathways such as neurodegeneration and autophagy, which are well-documented in cancer treatment and linked to mTOR signaling [48, 49]. Additionally, newly predicted diseases included breast cancer (BRCA) and lung adenocarcinoma (LUAD), both supported by preclinical evidence [46, 50].

**Fig. 14.**
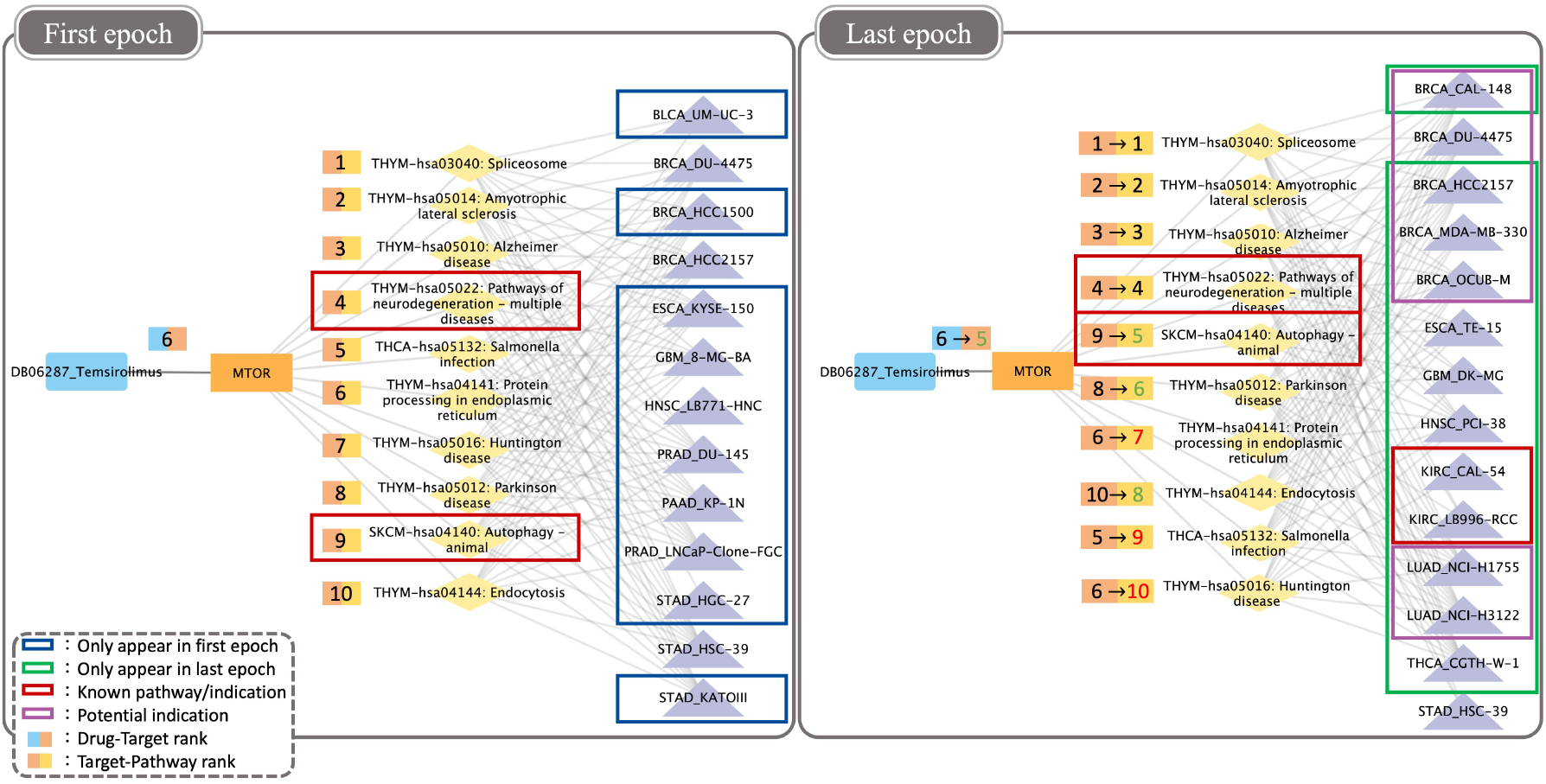
Case Study 2: D-T-P-I Networks of Temsirolimus.

These results demonstrate that GTCN is capable of not only maintaining the relevance of initially recognized drug indications and pathways but also recovering additional known treatment relationships while uncovering potential new therapeutic applications. This highlights the model’s ability to refine biological network representations dynamically, integrating both prior knowledge and new insights into drug-disease relationships, emphasizing its potential for identifying drug repurposing opportunities.

## Conclusion

In this paper, we proposed Graph Transformer-Convolution Networks (GTCNs) for drug-cancer association prediction, integrating heterogeneous biological networks with advanced encoding methods to capture drug-target-pathway-disease (D-T-P-I) relationships. Our model addresses two key challenges: inefficient utilization of biological networks and lack of interpretability. By incorporating biologically meaningful encodings for drugs, targets, pathways, and diseases, GTCNs enhance the understanding of drug mechanisms. Additionally, Graph Transformer Networks dynamically refine biological networks, filtering noise while emphasizing critical relationships, leading to improved predictive accuracy. Furthermore, attention-based interpretability provides insights into how drugs influence targets and regulate pathways, offering valuable explanations for the model’s decisions.

Experimental results on the REDDA benchmark and our cell line-based dataset demonstrated GTCNs’ superior performance, particularly in identifying positive drug-disease associations despite class imbalance. Our interpretability framework also revealed new therapeutic insights, including potential repurposing of methotrexate and temsirolimus for additional cancers, supported by literature evidence.

Despite these promising results, GTCNs have limitations that future work aims to address. Our current dataset scope could be expanded to include a broader range of drug types (e.g., protein-based drugs), disease categories, and additional biological nodes such as side effects. Furthermore, incorporating more advanced heterogeneous graph network models may enhance the framework’s ability to model complex interactions. Additionally, transitioning from binary association predictions to bioactivity prediction, such as IC50 values, would improve the model’s practical applicability in drug discovery and repurposing. These improvements will further enhance GTCNs’ versatility in computational pharmacology and precision medicine.

## Key Contributions

- **Comprehensive biological network encoding:** Unlike existing methods that primarily encode target nodes (e.g., drugs and diseases), we extend biologically meaningful encodings to all node types, including targets and pathways, improving feature representation and capturing complex biological interactions.
- **Dynamic graph refinement through GTNs:** We leverage Graph Transformer Networks (GTNs) to dynamically refine noisy and incomplete biological networks, enhancing the predictive power of the model by emphasizing critical relationships while filtering less relevant connections.
- **Enhanced interpretability in drug repurposing:** Our attention-based interpretability framework enables detailed analysis of model decisions, providing insights into drug-target-pathway-disease (D-T-P-I) relationships and identifying biologically plausible mechanisms for drug repurposing.

## Notes

### Competing Interest Statement

The authors have declared no competing interest.

## References

1. Nicola Nosengo. New tricks for old drugs. Nature, 534:314–316, 2016.

2. Jack W. Scannell, Alex Blanckley, Helen Boldon, and Brian Warrington. Diagnosing the decline in pharmaceutical r&d efficiency. Nature Reviews Drug Discovery, 11:191–200, 2012.

3. Sudeep Pushpakom, Francesco Iorio, Patrick A. Eyers, and et al. Drug repurposing: progress, challenges and recommendations. Nature Reviews Drug Discovery, 18:41–58, 2019.

4. J. G. Moffat, F. Vincent, J. A. Lee, J. Eder, and M. Prunotto. Opportunities and challenges in phenotypic drug discovery: an industry perspective. Nature Reviews Drug Discovery, 16:531–543, 2017.

5. J. Eder, R. Sedrani, and C. Wiesmann. The discovery of first-in-class drugs: origins and evolution. Nature Reviews Drug Discovery, 13:577–587, 2014.

6. N. Greives and H. X. Zhou. Both protein dynamics and ligand concentration can shift the binding mechanism between conformational selection and induced fit. Proceedings of the National Academy of Sciences USA, 111:10197–10202, 2014.

7. L. Pinzi and G. Rastelli. Molecular docking: shifting paradigms in drug discovery. International Journal of Molecular Sciences, 20, 2019.

8. X. Yang, Y. F. Wang, R. Byrne, G. Schneider, and S. Y. Yang. Concepts of artificial intelligence for computer-assisted drug discovery. Chemical Reviews, 119:10520–10594, 2019.

9. D. C. Swinney and J. Anthony. How were new medicines discovered? Nature Reviews Drug Discovery, 10:507–519, 2011.

10. J. L. Medina-Franco, M. A. Giulianotti, G. S. Welmaker, and R. A. Houghten. Shifting from the single to the multitarget paradigm in drug discovery. Drug Discovery Today, 18:495–501, 2013.

11. F. Vincent. et al. Phenotypic drug discovery: recent successes, lessons learned and new directions. Nature Reviews Drug Discovery, 21:899–914, 2022.

12. Yurui Chen and Louxin Zhang. How much can deep learning improve prediction of the responses to drugs in cancer cell lines? Briefings in Bioinformatics, 23:1–8, 2022.

13. Mei Li, Xiangrui Cai, Sihan Xu, and Hua Ji. Metapath-aggregated heterogeneous graph neural network for drug–target interaction prediction. Briefings in Bioinformatics, 24(1):bbac578, 01 2023.

14. Yuxuan Gu, Shuang Zheng, Qian Yin, Rong Jiang, and Jinhui Li. Redda: Integrating multiple biological relations to heterogeneous graph neural network for drug-disease association prediction. Computers in Biology and Medicine, 150:106127, 2022.

15. Seongjun Yun, Minbyul Jeong, Raehyun Kim, Jaewoo Kang, and Hyunwoo J. Kim. Graph transformer networks, 2020.

16. Seongjun Yun, Minbyul Jeong, Sungdong Yoo, Seunghun Lee, Sean S. Yi, Raehyun Kim, Jaewoo Kang, and Hyunwoo J. Kim. Graph transformer networks: Learning meta-path graphs to improve gnns. Neural Networks, 153:104–119, 2022.

17. Thomas N. Kipf and Max Welling. Semi-supervised classification with graph convolutional networks, 2017.

18. C. Knox, M. Wilson, C. M. Klinger, M. Franklin, E. Oler, A. Wilson, A. Pon, J. Cox, N. E. L. Chin, S. A. Strawbridge, M. Garcia-Patino, R. Kruger, A. Sivakumaran, S. Sanford, R. Doshi, N. Khetarpal, O. Fatokun, D. Doucet, A. Zubkowski, D. Y. Rayat, H. Jackson, K. Harford, A. Anjum, M. Zakir, F. Wang, S. Tian, B. Lee, J. Liigand, H. Peters, R. Q. R. Wang, T. Nguyen, D. So, M. Sharp, R. da Silva, C. Gabriel, J. Scantlebury, M. Jasinski, D. Ackerman, T. Jewison, T. Sajed, V. Gautam, and D. S. Wishart. Drugbank 6.0: the drugbank knowledgebase for 2024. Nucleic Acids Res., 52(D1):D1265–D1275, 2024.

19. Cancer Genome Atlas Research Network, J. N. Weinstein, E. A. Collisson, G. B. Mills, K. R. Shaw, B. A. Ozenberger, K. Ellrott, I. Shmulevich, C. Sander, and J. M. Stuart. The cancer genome atlas pan-cancer analysis project. Nat Genet., 45(10):1113–1120, 2013.

20. M. Kanehisa and S. Goto. Kegg: kyoto encyclopedia of genes and genomes. Nucleic Acids Res., 28(1):27–30, 2000.

21. W. Yang, J. Soares, P. Greninger, E. J. Edelman, H. Lightfoot, S. Forbes, N. Bindal, D. Beare, J. A. Smith, I. R. Thompson, S. Ramaswamy, P. A. Futreal, D. A. Haber, M. R. Stratton, C. Benes, U. McDermott, and M. J. Garnett. Genomics of drug sensitivity in cancer (gdsc): a resource for therapeutic biomarker discovery in cancer cells. Nucleic Acids Res., 41(Database issue):D955–D961, 2013.

22. N. Cheng. et al. Hansynergy: Heterogeneous graph attention network for drug synergy prediction. J Chem Inf Model, 64(10):4334–4347, 2024.

23. J. Lee. et al. Dlm-dti: a dual language model for the prediction of drug-target interaction with hint-based learning. J Cheminform, 16(1):14, 2024.

24. C. Y. Lin, C. H. Lee, Y. H. Chuang, J. Y. Lee, Y. Y. Chiu, Y. H. Wu Lee, Y. J. Jong, J. K. Hwang, S. H. Huang, L. C. Chen, C. H. Wu, S. H. Tu, Y. S. Ho, and J. M. Yang. Membrane protein-regulated networks across human cancers. Nat Commun., 10(1):3131, 2019.

25. A. P. Lind and P. C. Anderson. Predicting drug activity against cancer cells by random forest models based on minimal genomic information and chemical properties. PLoS One, 14(7):e0219774, 2019.

26. D. Szklarczyk, R. Kirsch, M. Koutrouli, K. Nastou, F. Mehryary, R. Hachilif, A. L. Gable, T. Fang, N. T. Doncheva, S. Pyysalo, P. Bork, L. J. Jensen, and C. von Mering. The string database in 2023: protein-protein association networks and functional enrichment analyses for any sequenced genome of interest. Nucleic Acids Res., 51(D1):D638–D646, 2023.

27. Ning Cheng, Li Wang, Yiping Liu, Bosheng Song, and Changsong Ding. Hansynergy: Heterogeneous graph attention network for drug synergy prediction. Journal of Chemical Information and Modeling, 64(10):4334–4347, 2024. PMID: 38709204.

28. Jonghyun Lee, Dae Won Jun, Ildae Song, and Yun Kim. Dlm-dti: a dual language model for the prediction of drug-target interaction with hint-based learning. JOURNAL OF CHEMINFORMATICS, 16(1), FEB 1 2024.

29. N. Haider. Functionality pattern matching as an efficient complementary structure/reaction search tool: an open-source approach. Molecules, 15(8):5079–5092, 2010.

30. S. Kim, J. Chen, T. Cheng, A. Gindulyte, J. He, S. He, Q. Li, B. A. Shoemaker, P. A. Thiessen, B. Yu, and et al. Pubchem in 2021: new data content and improved web interfaces. Nucleic Acids Research, 49(D1):D1388–D1395, 2021.

31. R. D. Taylor, M. MacCoss, and A. D. Lawson. Rings in drugs: Miniperspective. Journal of Medicinal Chemistry, 57(14):5845–5859, 2014.

32. Zhouxin Yu, Feng Huang, Xiaohan Zhao, Wenjie Xiao, and Wen Zhang. Predicting drug–disease associations through layer attention graph convolutional network. Briefings in Bioinformatics, 22(4):bbaa243, 10 2020.

33. Wei Zhang, Xuekai Yue, Yujie Chen, Fang Huang, and Feng Liu. Scmfdd: Similarity constrained matrix factorization with dual consistency for predicting drug-disease associations. Bioinformatics, 34(2):315–324, 2018.

34. Wenqing Wang, Sheng Yang, Xiantao Zhang, and Jiajie Li. Drug repositioning by integrating target information through a heterogeneous network model. Bioinformatics, 30(20):2923–2930, 2014.

35. Xiaofeng Chen, Chengjun Yan, Xiantao Zhang, and Zhi-Hong You. A drug–disease association prediction method based on a novel similarity kernel fusion strategy. Bioinformatics, 31(12):2001–2008, 2015.

36. Bo-Wei Zhao, Lun Hu, Zhu-Hong You, Lei Wang, and Xiao-Rui Su. Hingrl: predicting drug–disease associations with graph representation learning on heterogeneous information networks. Briefings in Bioinformatics, 23(1):bbab515, 12 2021.

37. Jiezhong Ma, Simon Fong, Rebecca Wong, et al. Lagcn: Layered attention graph convolutional networks for drug repurposing. BMC Bioinformatics, 22(1):34, 2021.

38. Yajie Meng, Changcheng Lu, Min Jin, Junlin Xu, Xiangxiang Zeng, and Jialiang Yang. A weighted bilinear neural collaborative filtering approach for drug repositioning. Briefings in Bioinformatics, 23(2):bbab581, 01 2022.

39. R Sehrawat, P Rathee, S Khatkar, E Akkol, M Khayatkashani, S. M Nabavi, and A Khatkar. Dihydrofolate reductase (dhfr) inhibitors: A comprehensive review. Current Medicinal Chemistry, March 2023. Epub ahead of print.

40. Early Breast Cancer Trialists’ Collaborative Group (EBCTCG). Long-term outcomes for neoadjuvant versus adjuvant chemotherapy in early breast cancer: meta-analysis of individual patient data from ten randomised trials. Lancet Oncol, 19(1):27–39, 2018.

41. J. Chang, H. Wu, and J. Wu. et al. Constructing a novel mitochondrial-related gene signature for evaluating the tumor immune microenvironment and predicting survival in stomach adenocarcinoma. J Transl Med, 21(1):191, 2023.

42. L. Chen and Z. Zhang. et al. Nadph production by the oxidative pentose-phosphate pathway supports folate metabolism. Nat Metab, 1:404–415, 2019.

43. E.E.W. Cohen. et al. Pembrolizumab versus methotrexate, docetaxel, or cetuximab for recurrent or metastatic head-and-neck squamous cell carcinoma (keynote-040). Lancet, 393:156–167, 2019.

44. P.M. Wilson. et al. Standing the test of time: targeting thymidylate biosynthesis in cancer therapy. Nat Rev Clin Oncol, 11:282–298, 2014.

45. E. Niemela. et al. Nanoparticles carrying fingolimod and methotrexate enables targeted induction of apoptosis and immobilization of invasive thyroid cancer. Eur J Pharm Biopharm, 148:1–9, 2020.

46. D. Miricescu. et al. Pi3k/akt/mtor signaling pathway in breast cancer: From molecular landscape to clinical aspects. Int J Mol Sci, 22(1):173, 2020.

47. J. Hsieh. et al. Renal cell carcinoma. Nat Rev Dis Primers, 3:17009, 2017.

48. G.Y. Liu. et al. mtor at the nexus of nutrition, growth, ageing and disease. Nat Rev Mol Cell Biol, 21(4):183–203, 2020.

49. N. Deleyto-Seldas and A. Efeyan. The mtor-autophagy axis and the control of metabolism. Front Cell Dev Biol, 9:655731, 2021.

50. H.W. Chang. et al. Therapeutic effect of repurposed temsirolimus in lung adenocarcinoma model. Front Pharmacol, 9:778, 2018.

